# Efficiency of heterogenous functional connectomes explains variance in callous-unemotional traits after computational lesioning of cortical midline and salience regions

**DOI:** 10.1101/2022.10.07.511379

**Authors:** Drew E. Winters, Daniel R. Leopold, Joseph T. Sakai, R. McKell Carter

## Abstract

Callous-unemotional (CU) traits are a youth antisocial phenotype hypothesized to be a result of differences in the integration of multiple brain systems. However, mechanistic insights into these brain systems are a continued challenge. Where prior work describes activation and connectivity of the connectome in relation to these systems, new mechanistic insights can be derived by removing nodes and characterizing changes in network properties (hereafter referred to as computational lesioning) to characterize the resilience and vulnerability of the brain’s functional connectome. Here, we study the resilience of connectome integration in CU traits by estimating changes in efficiency after computationally lesioning individual-level connectomes. From resting-state data of 86 participants (48% female, age 14.52±1.31) drawn from the Nathan Kline institute’s Rockland study, individual-level connectomes were estimated using graphical lasso. Computational lesioning was conducted both sequentially and by targeting global and local hubs. We calculated changes in network efficiency after each lesion. Then, elastic net regression was applied to determine how these changes explained variance in CU traits. Follow-up analyses characterized modeled node hubs, examined moderation, determined impact of targeting, and decoded the brain mask by comparing regions to meta-analytic maps. Elastic net regression revealed that computational lesioning of 23 nodes, network modularity, and Tanner stage explained variance in CU traits. Hub assignment of selected hubs differed at higher CU traits. No evidence for moderation between simulated lesioning and CU traits was found. Targeting global hubs increased efficiency and targeting local hubs had no effect at higher CU traits. Identified brain mask meta-analytically associated with more emotion and cognitive terms. Although reliable patterns were found across participants, adolescent brains were heterogeneous even for those with a similar CU traits score. Adolescent brain response to simulated lesioning revealed a pattern of connectome resiliency and vulnerability that explains variance in CU traits, which can aid prediction of youth at greater risk for higher CU traits.

---

Callous-Unemotional (CU) traits are an antisocial phenotype in youth related to the affective impairments in adult psychopathy (Barry et al., 2000; Frick et al., 2014a; Frick & White, 2008) and are defined by impairments in prosocial emotions such as remorse, guilt, and empathy (Frick et al., 2014a, 2014b). CU traits associate with persistent criminal behavior (Kahn et al., 2013) and higher substance use (Winters et al., 2020) while causing substantial costs to society (Kiehl & Hoffman, 2011). Available treatments, however, are limited in their efficacy, indicating the need to better understand mechanisms underlying these traits (for review: White et al., 2022). Brain mechanisms underlying CU traits have been conceptualized as involving either under activation of modular regions involved in salience (Blair & Frith, 2000; Patrick, 1994) or less efficient integration between multiple brain networks (impaired integration theory; Hamilton et al., 2015). But, extracting mechanisms from this work is an ongoing challenge, which is plausibly due to the following three reasons. First, rather than modular activation, to understand integration across multiple brain systems, we need to consider how different and potentially distant hubs respond to perturbations of a given network node (i.e., node removal) using computationally simulated lesions (Honey & Sporns, 2008). Second, the shape of brain networks and resulting information processing streams (i.e., topology; De Vico Fallani et al., 2014; Kaiser et al., 2015) are rarely examined in studies on CU traits. Finally, the substantial heterogeneity of individual connectomes (Damoiseaux et al., 2021) and their heterogeneous association with both psychopathy (Dotterer et al., 2020) and CU traits (Winters, Sakai, et al., 2021) need accounted for to make accurate inferences (Gates & Molenaar, 2012; Molenaar, 2004). Thus, the present study examines topological brain properties underlying CU traits by examining individual-level brain responses with computational lesioning to understand the functional architecture accounting for variance in CU traits and underlying etiology.

Etiological theories of CU traits posit primary impairments that center on either affective (Blair, 2008; Hawes & Dadds, 2012) or cognitive deficits (i.e., attention and cognitive control; Hamilton & Newman, 2018) with both positions supported by neurobiological evidence. For example, affective processing deficits include salience regions such as the insula and amygdala (Seara-Cardoso et al., 2016) with the majority of studies focusing on the amygdala in psychopathy (Blair & Frith, 2000; Patrick, 1994) and CU traits (Marsh et al., 2008). Cognitive deficits involve social and control regions in cortical midline structures such as the medial/lateral prefrontal cortices and anterior/posterior cingulate. Less activation of these regions are observed during top-down attention (Larson et al., 2013; Newman & Baskin-Sommers, 2012), reward anticipation (Veroude et al., 2016), and decision making tasks involving conflict monitoring (Abe et al., 2018; White et al., 2013). Whether these cognitive results indicate a general attention impairment or a tendency to not process affective information in the present context is debated (Blair & Mitchell, 2009); however, the ability to monitor and bring attention to context for regulating goal-directed behaviors (i.e., cognitive control; Botvinick et al., 2001) is an important impairment associated with CU traits (e.g., Gluckman et al., 2016) and related to decrements in representing others’ affective states (e.g., Winters & Sakai, 2021) that associate with differences in these multiple brain systems (e.g., Winters et al., 2022).

Impaired integration across these multiple brain systems (e.g., control, social, salience) are thought to underlie CU trait impairments beyond modular activation (impaired integration hypothesis; Hamilton et al., 2015). Studies supporting this demonstrate less integration within and between networks involving regions outlined above including the default mode (DMN), frontoparietal (FPN), and salience (SAL) networks. For example, where we would expect greater connectivity within networks among healthy brains, CU traits associate with less connectivity within the DMN (Cohn et al., 2015; Umbach & Tottenham, 2020), FPN (Winters, Sakai, et al., 2021), and SAL (Yoder et al., 2016). Functional integration of these networks also associates with normative brain processes (e.g., DMN and cognitive empathy: Winters, Pruitt, et al., 2021) that are perturbed among individuals with elevated CU traits. Furthermore, where we would expect an anticorrelation between task positive and task negative networks in typically developing brains (Uddin et al., 2009), higher CU traits associate with a diminished anticorrelation between the DMN and SAL (Winters & Hyde, 2022), as well as between the DMN and FPN (Pu et al., 2017; Winters, Sakai, et al., 2021), which has also been found in adult psychopathy (Dotterer et al., 2020). This pattern of less connectivity within and between these networks is theorized to underlie cognitive impairments that impact affective processing (Hamilton et al., 2015) possibly via difficulties with perspective taking and cognitive control (Winters et al., 2022).

It is necessary to also highlight the lack of convergence of the brain literature on CU traits and psychopathy that appears to point to more general topological properties of the brain. For example, task-based findings demonstrate some overlap but broadly heterogenous activation patterns among similar tasks (e.g., Seara-Cardoso et al., 2022); some connectivity studies did not find aberrant connectivity in the DMN (Pu et al., 2017) or in the SAL and FPN (Umbach & Tottenham, 2020) whereas other studies reviewed above did. One possibility is that the brains of those higher in CU traits may have a less modular structure. Modularity describes the structure of the brain network representing the strength of divisions into network modules with more modularity indicating dense connections within the network and less connections between networks (Newman & Girvan, 2004). Less modular structure of the brain results in less efficiency (Tosh & McNally, 2015), which impacts behavior and cognitive functioning (Rypma & Prabhakaran, 2009). Decreased modularity could manifest in different networks of the brain (e.g., less connectivity of the DMN or the SAL) that could plausibly represent different subgroups of CU trait phenotypes. For example, CU traits have multiple presentations underlying individual differences (Fanti et al., 2013; Fanti et al., 2018; Hadjicharalambous & Fanti, 2018; Sebastian et al., 2012) and it is plausible that differential decrements in modularity of the DMN compared to the SAL may underly these different subgroups. These hypotheses remain speculative, however, and substantial methodological improvements are needed in this area of study.

The following three methodological improvements can help improve the characterization of neural substrates of CU traits. *<u>First</u>*, computational lesioning has yet to be applied to understand underlying functional brain properties of youth with CU traits. Computational lesioning probes the resilience or vulnerability of a functional network related to specific nodes, thereby providing valuable insights into entry points for mechanistic understanding of a disorder (Deco & Kringelbach, 2014). Although real lesions involve long-term brain re-organization that is not captured when simulating lesions, computational lesioning confirms far reaching disruptions of functional architecture that are related to the modular structure of nodes (e.g., global versus local hubs) that is found in real lesions (Gratton et al., 2012; Tao & Rapp, 2021). Specifically, characterizing the disruptions that these different hubs cause on the connectome after computational lesioning are important for understanding underlying brain function (Honey & Sporns, 2008).

*<u>Second</u>*, although rarely studied in CU trait investigations, considering topological properties of the brain such as efficiency, modularity, and hubs can characterize how information is transferred in the brain. For example, measuring *efficiency* captures how information is exchanged in a network assuming that shorter distances between nodes is more efficient (Achard & Bullmore, 2007; Latora & Marchiori, 2001; Rubinov & Sporns, 2010), *modularity* captures within module density (Newman & Girvan, 2004), and *hubs* that are global or local captures weather a node is more connected across the brain or within module (Gratton et al., 2012; Tao & Rapp, 2021). The few studies of CU traits examining topology have identified important differences (Dotterer et al., 2020; Jiang et al., 2021; Winters, Sakai, et al., 2021) that go beyond the typical considerations of functional activation and connection strength.

*<u>Third</u>*, letting go of the incorrect assumption of homogeneity in brain analyses can improve our statical inferences on potential mechanisms. For example, most relevant work uses group averages across the brain to make inferences. This would suggest an assumption of strict homogeneity of individual brains (Gates, 2022), but it is well known that functional brain patterns are as unique as fingerprints (Damoiseaux et al., 2021), thus the assumption of homogeneity incorrect. Accordingly, recent studies demonstrate substantial heterogeneity of the brain in relation to CU traits (Winters, Sakai, et al., 2021) and psychopathy (Dotterer et al., 2020). These studies demonstrate some shared patterns despite this heterogeneity (known as weak homogeneity; Gates, 2022), which appears to more accurately reflect how we should model the brain. Failing to account for this individual variability when examining patterns across individuals leads to inaccurate inferences (Gates & Molenaar, 2012; Molenaar, 2004). Thus, examining connectome efficiency across individual-level connectives using computational lesioning can improve our mechanistic understanding of CU traits by probing the resilience and vulnerability of an individuals’ functional connectome.

The way nodes in the functional connectome are connected can have different impacts on efficiency after computational lesioning. For example, a node can be globally connected, meaning it has denser connections across the brain between network modules, or can be locally connected, meaning it has denser connections within its respective network module and less connections with other modules – these are called global and local hubs, respectively (Gratton et al., 2012; Tao & Rapp, 2021). Computational lesioning of global hubs reduces the number of long-range connections, resulting in more segregation of network modules, whereas lesioning of local hubs reduces the number of short-range connections, resulting in less segregation of network modules (Gratton et al., 2012; Sporns et al., 2007; Tao & Rapp, 2021). These forms of reorganization of the brain’s modularity can have substantial impact on the brain’s efficiency (e.g., Tosh & McNally, 2015). Efficiency in the brains of those with CU traits is lower than controls (Jiang et al., 2021), which may be related to the global or local hub-like qualities (i.e., “hubness”) of a given node. Despite not yet having been investigated, to our knowledge, data on how the efficiency of a network changes related to properties of a node being a global or local hub could provide practical, functionally important mechanistic insights (Deco & Kringelbach, 2014). Thus, an important next step for understanding mechanisms underlying CU traits is to examine changes in efficiency using computational lesioning, characterize nodal “hubness”, and examine targeted lesioning of node types in relation to CU traits.

The present study examines changes in efficiency after computationally simulating lesions in relation to CU traits among a community sample of adolescents. CU traits exist along a continuum in community samples with substantial evidence of similar neurocognitive and neurobiological impairments as forensic samples with these traits (Seara-Cardoso et al., 2022; Viding & McCrory, 2012). In this sample of community adolescents, we hypothesize that changes in efficiency after computationally simulating lesions can explain variance in and help us understand mechanisms underlying CU traits. Specifically, we hypothesize that changes in response to cortical midline structures as well as regions associated with salience (amygdala and insula) will account for variance in CU traits. Consistent with the literature on global and local hub lesioning discussed above, we specifically anticipate that lesioning local hubs will demonstrate the greatest decrement in efficiency. Additionally, in accordance with prior studies, we expect modularity will account for variance in CU traits such that lower modularity will associate with higher CU traits. This information has the potential to provide unique mechanistic insights that complement task-based and functional connectivity studies by identifying points of resilience and vulnerability in functional connectomes in relation to CU traits.

## Methods

### Sample

Participants were drawn from the Rockland study collected by the Nathan Kline Institute. We downloaded raw fMRI files and study measurements from the 1000 connectomes project (www.nitrc.org/projects/fcon_1000/). We included adolescent participants between the ages of 13-17 with an IQ > 80 as measured by the WAIS-II (α = .96; Wechsler, 2011). We excluded potential participants with motion > 3mm or > 20% of invalid fMRI scans. Two participants had spikes in motion that was near the end of the session and were able to retain those participants by cutting their time series leaving a total analysis sample of 86. The participants in this sample were predominantly White (White= 63%, Black = 24%, Asian = 9%, Indian = 1%, other= 3%) with 14% reporting Latinx ethnicity, mean age of 14.5 (14.52±1.31) years, slightly more males (females = 48%), and mean pubertal development just below full maturity (4.10±0.97; range 1-5). Nooner et al. (2012) outlines study procedures, including consent and assent for all participants.

### Measures

#### Inventory of Callous-Unemotional Traits (ICU)

CU traits were assessed using the total score of the 24-item ICU (Frick, 2004). Consistent with Kimonis et al. (2008), we removed two items with poor psychometrics, which had adequate reliability in the current sample (α=0.72). Higher scores indicate greater CU traits. The total ICU score was used as the primary outcome of interest.

#### Tanner Stage

Sex and pubertal stage were measured using the Tanner assessment (α = 0.77). Parents rated pictures of secondary sex characteristics indicating pubertal development of 1 (pre-pubertal) to 5 (full maturity; Petersen et al., 1988).

#### fMRI Acquisition

Resting state fMRI images from the Rockland dataset were collected by the Nathan Kline Institute using a Siemens TimTrio 3T scanner with a blood oxygen level dependent (BOLD) contrast and an interleaved multiband echo planar imaging (EPI) sequence. Each scan involved resting state (260 EPI volumes; repetition time (TR) 1400ms; echo time (TE) 30ms; flip angle 65°; 64 slices, Field of view (FOV) = 224mm, voxel size 2mm isotropic, duration = 10 minutes) and a magnetization prepared rapid gradient echo (MPRAGE) anatomical image (TR= 1900ms, flip angle 9°, 176 slices, FOV= 250mm, voxel size= 1mm isotropic). The Siemens sequence does not collect images until T1 stabilization is achieved, so removing scans was not necessary. Instructions for participants were to keep their eyes closed without falling asleep and to not think of anything while they let their mind wander.

#### Resting-State fMRI Preprocessing

Imaging data was preprocessed with the standard preprocessing in the CONN toolbox (version 18b; Whitfield-Gabrieli & Nieto-Castanon, 2012) that uses Statistical Parametric Mapping (SPM version 12; Penny et al., 2011). Motion outliers were flagged for correction if > 0.5mm using the Artifact Detection Tools (ART; http://www.nitrc.org/projects/artifact_detect) and regressed out using spike regression. Slice timing correction was not used given the fast multiband acquisition (Glasser et al., 2013; Wu et al., 2011). The anatomic component-based noise correction method (aCompCor; Whitfield-Gabrieli & Nieto-Castanon, 2012) was used to regress out CSF and white matter noise. MPRAGE and EPI images were co-registered and normalized to an MNI template; and data was bandpass filtered between 0.008 and ,09HZ to retain resting state signals. Finally, we parcellated this data into 164 ROIs using the Harvard Oxford atlas for cortical and sub-cortical areas (Desikan et al., 2006) as well as the Automated Anatomical Labeling Atlas for cerebellar areas (Tzourio-Mazoyer et al., 2002).

We found 24 participants had excess motion > 3mm and four had >20% of invalid scans. However, we were able to retain two of the participants with excess motion because this motion was at the end of the timeseries and was able to be snipped while still retaining > 90% of the timeseries. This left a total of 86 participants for analysis.

### Construction of Individual-Level Functional Connectomes

Single-subject connectomes were derived within python (version 3.9.5; Van Rossum G. & L., 2009; code for all anlayses can be found here: https://doi.org/10.17605/OSF.IO/M5G6X) using the graphical lasso covariance estimator in the package “scikit-learn” (Pedregosa et al., 2011). This resulted in a brain wide sparse precision matrix for each participant. As opposed to imposing arbitrary covariance thresholds that reflect unique characteristics of the sample, this sparse matrix approach was chosen because it uses a principled approach that retains meaningful connections after conditioning on the rest of the matrix (Smith et al., 2011; Varoquaux et al., 2010). Whole brain ROIs were defined using the Harvard Oxford atlas for cortical and sub-cortical areas (Desikan et al., 2006) as well as the Automated Anatomical Labeling Atlas for cerebellar areas (Tzourio-Mazoyer et al., 2002).

### Individual-Level Brain Connectivity Measures

Functional brain properties were derived from network estimations using the python-based brain connectivity toolbox (Rubinov & Sporns, 2010). First, we applied the robust Louvain algorithm (Blondel et al., 2008; Lancichinetti & Fortunato, 2009) combined with an iterated fine tuning algorithm to optimize identifying modular structures at the single-subject level (Sun et al., 2009) and address the Louvain algorithm’s stochastic nature (Bassett et al., 2011). Specifically, we estimated five communities for each participant to determine the optimal individual-level gamma parameter, re-estimated communities using the individual-level optimal gamma, calculated the similarity for each iteration for each participant, then derived a single consensus community across all iterations at the single-subject level.

Using the tuned parameters from above, we calculated individual-level modularity, participation coefficient, and within-module degree z-score. The participation coefficient was used to identify global hubs because it represents the strength of cross module connections; conversely the within module z-score was used to identify local hubs because it measures the extent of intra-modular connections of each node (Guimera & Nunes Amaral, 2005). To identify each the hubness of each node we used criteria by Tao and Rapp (2021) but, where they used mean and standard deviation, we used median and median absolute deviation (MAD) because it is not as subject to sampling effects (Leys et al., 2013). To identify individual-level global hubs (participation coefficient > 1 MAD + median) and local hubs (within-module degree z-score coefficient > 1 MAD + median). Consistent with prior computational lesion studies (He et al., 2009), we identified non-hubs that were connector (i.e., more likely to be connected across network) or periphery nodes (i.e., more connected within module) but did not meet hub criteria. Finally, we calculated individual baseline efficiency to compare changes in efficiency after computationally lesioning.

### Computationally Simulated Lesions at the Individual-Level

To investigate the dynamics and probe resilience or vulnerability underlying individual-level connectomes, we applied a procedure to computationally simulate lesions (deletion) over the functional connectome (e.g., He et al., 2009). Specifically, we used node deletion across each participant’s functional connectome with two separate approaches. The first was a sequential deletion procedure for each node across the brain, and the second targeted nodes with properties of global and local hubs separately and connector and periphery non-hubs separately. After each computationally simulated lesion we calculated brain efficiency and subtracted each participant’s baseline efficiency score to assess the brain’s response.

### Feature Selection and Elastic Net Regression

Because adding all 164 nodes in one model would lead to noise and/or spurious associations, we conducted feature selection to retain only those features that are the most pertinent and improve model performance (Dosenbach et al., 2010). Specifically, we used the k-best feature selection using the f regression function within “scikit-learn” (Pedregosa et al., 2011) and conducted hyperparameter tuning of the number of k features selected to improve model performance. This resulted in 23 nodes (23 features) representing changes in efficiency for each participant’s functional connectome after removing that node, (24^th^ feature) overall brain modularity, and (25^th^ feature) Tanner stage (see results). We also included (26^th^ feature) sex independently of feature selection because of its implications for differences in the brain and CU traits in youth (Raschle et al., 2018). Importantly, head motion was not selected as accounting for substantial variance thus, was not included in the model. Brain features were placed in the model as independent variables to decode what brain activity predicts CU traits.

A linear elastic net regression was implemented in the python package “scikit-learn” (Varoquaux et al., 2010) to evaluate the relationship between functional connectome change in efficiency after removing each node. Model performance was evaluated using a nested five-fold cross-validation procedure involving hyper-parameter tuning and cross-validation. We used mean squared error, R^2^, and mean absolute error to evaluate the model as well as comparing training and testing cross-validation scores. Additionally, we compared the model to a dummy model to assess if the model performed better than chance using the mean squared error. We then assessed cumulative empirical distribution of model performance under the null hypothesis using a permutation test with 2000 iterations. Results were considered statistically significant if 95% of these 2000 R^2^ values were lower than the R^2^ of the real data (Dosenbach et al., 2010).

### Meta-Analytic Decoding

For connections response to brain regions accounting for variation in CU traits, we used Neurosynth (http://neurosynth.org) to meta-analytically annotate their functional characteristics. The automated neuroimaging meta-analysis computed whole-brain posterior activation distributions *P(Term | Activation)* for the psychological concepts examined (Yarkoni et al., 2011). Terms that have been consistently associated with a particular activation map can be identified using unbiased reverse-inference analyses across the Neurosynth database. At the time of writing this paper, Neurosynth database contained maps for 1,334 terms, 507,891 coordinates extracted from 14,371 fMRI studies, and coactivation maps for 150,000 brain locations. Using this database, we first decoded the mask of regions in our analysis, and second, identified regional terms loading on to CU traits and psychopathy.

First, to decode the mask of identified ROIs during feature selection, we took the coordinates of all 23 regions and created a mask with 8mm spheres around the center of each coordinate using the python package ‘nltools’ (Chang, 2020) that is publicly available (https://doi.org/10.17605/OSF.IO/M5G6X and https://identifiers.org/neurovault.collection:12738). This mask was used to then characterize co-activation of brain regions across studies in the Neurosynth database. Statistical inference for each voxel in the brain volume was conducted using Chi-Square tests to generate a z-value map thresholded with a false discover-rate adjusted p-value of p< 0.01; and the voxel-wise correlation coefficient between the co-activation map and each term-specific map was assessed for extraction. To obtain a large enough number to account for variation, the top 40 terms (e.g., executive functioning, affect response; excluding methodological techniques [e.g., fMRI] or anatomy terms [e.g., prefrontal cortex]) were taken as the most likely associated term. We assessed ambiguous terms by examining the top ten loading studies to determine inclusion or exclusion. For example, the term “amygdala response” involved the study of response to emotional stimuli, thus it was included given it is the brains response to emotional stimuli. We averaged r values for terms that have common root terms (e.g., feeling and feelings). Additionally, we placed each term in a general category (e.g., emotion, cognitive) that represents the studies under that term.

Second, we examined identified region terms and extracted loading values on CU traits and psychopathy. We choose CU traits and psychopathy as our primary terms because CU traits represent the affective dimension of psychopathy (Barry et al., 2000; Frick et al., 2014a) and brain features are shared between psychopathic adults and youth with CU traits (Seara-Cardoso et al., 2022). We searched variations of terms for CU traits (callous-unemotional, callous, callousness) and psychopathy (psychopathic, psychopath), reviewed the highest loading studies to ensure construct consistency, and extracted the loading value for that brain area and term.

### Random Effects Models, Node Characterization, and Evaluating Moderation

To contextualize brain properties related to the model results, linear random effects models were conducted in python (version 3.9.5; Van Rossum G. & L., 2009) using the “statsmodels” package (Seabold & Perktold, 2010) to characterize selected nodes, test for mediation, and test the impact of targeting global and local nodes. For all models, we accounted for random intercept variation for each participant. Confidence intervals for each parameter were bootstrapped with 2000 resamples.

We characterized nodes in the elastic net model by evaluating if their global or local “hubness” associated with CU traits. Importantly, nine nodes were excluded from analyses characterizing global hubs because zero participants had a global hub for that node. To assess whether nodes’ global or local “hubness” was different at higher CU traits from what was typically expected, we compared the probability of being a global or local hub for each node against a random distribution to assess if the current sample’s probability was better than chance.

We evaluated sex, modularity, and Tanner stage as potential moderators of above brain associations because both sex (Raschle et al., 2018) and pubertal stage (Cameron, 2004; Dahl, 2004; Sisk & Foster, 2004) demonstrate important differences in brain development and CU traits, and modularity can influence efficiency. First, we identified which of the above to evaluate based on significance in relation to CU traits in the elastic net model. We then ran correlations of the significant independent variables with every significant node in the elastic net regression and identified which nodes they may be a moderator for by selecting correlation values that were 2 median absolute deviations greater or less the median r value. Potential moderation terms were derived using the residualized centering approach using the python package ‘resmod’ (Winters, 2022). This approach orthogonalizes included terms by centering the residuals, thereby avoiding the violation of model assumptions by removing correlated residuals (Little et al., 2006). Confidence intervals were bootstrapped with 2000 resamples to test moderation.

## Results

### Lesioning Nodes Negatively Impacted Efficiency Across All Participants

Changes in efficiency after node deletion were, on average, negative for all nodes across the sample. The magnitude of this decrement varied by node (Table 1). Thus, positive associations in the following analyses represent the connectomes resilience because the change is closer to zero and negative associations represent a greater decrement in efficiency.

**Table 1.** Selected regions coordinates, abbreviations used, and descriptives on change in efficiency

| Area | Abbreviation | Coordinate |  |  | Efficiency Change |  |
| --- | --- | --- | --- | --- | --- | --- |
|  |  | X | Y | Z | Mean | SD |
| Cerebellar Anterior | Cebellar.Anterior | 0 | -63 | -30 | -0.0067 | 0.0004 |
| Inferior frontal gyrus R | Language.IFG R | 54 | 28 | 1 | -0.0065 | 0.0004 |
| Supramarginal gyrus L | Salience.SMG L | -60 | -39 | 31 | -0.0067 | 0.0005 |
| Anterior insula R | Salience.Ainsula R | 47 | 14 | 0 | -0.0067 | 0.0004 |
| Posterior Cingulate Cortex | DefalutMode.PCC | 1 | -61 | 38 | -0.0069 | 0.0005 |
| Vermis 9 | Ver9 | 1 | -55 | -35 | -0.0066 | 0.0005 |
| Vermis 3 | Ver3 | 1 | -40 | -11 | -0.0063 | 0.0004 |
| Amygdala L | Amygdala L | -23 | -5 | -18 | -0.0064 | 0.0004 |
| Planum Temporale R | PT R | 55 | -25 | 12 | -0.0067 | 0.0004 |
| Parietal Operculum L | PO L | -48 | -32 | 20 | -0.0065 | 0.0003 |
| Frontal Pole L | FP L | -40 | 18 | 5 | -0.0060 | 0.0004 |
| Temporal Fusiform Cortex Posterior Division R | pTFusC R | 36 | -24 | -28 | -0.0067 | 0.0004 |
| Temporal Fusiform Cortex Anterior Division L | aTFusC L | -32 | -4 | -42 | -0.0067 | 0.0004 |
| Cuneal L | Cuneal L | -8 | -80 | 27 | -0.0066 | 0.0004 |
| Anterior Cingulate | AC | 1 | 18 | 24 | -0.0067 | 0.0004 |
| Supplementary Motor Area L | SMA L | -5 | -3 | 56 | -0.0066 | 0.0004 |
| Intracalcarine Cortex L | ICC L | -10 | -75 | 8 | -0.0064 | 0.0004 |
| Superior Parietal Lobule L | SPL L | -29 | -49 | 57 | -0.0068 | 0.0004 |
| Postcentral Gyrus L | PostCG L | -38 | -28 | 52 | -0.0067 | 0.0004 |
| Inferior Temporal Gyrus Temporooccipital L | toITG L | -52 | -53 | -17 | -0.0069 | 0.0004 |
| Middle Temporal Gyrus Anterior Division L | aMTG L | -57 | -4 | -22 | -0.0069 | 0.0006 |
| Temporal Pole L | TP L | -40 | 11 | -30 | -0.0050 | 0.0008 |
| Inferior Frontal Gyrus Pars Opercularis L | IFG oper L | -51 | 15 | 15 | -0.0069 | 0.0005 |

Note: Efficiency change is the change in baseline efficiency after lesioning the brain region

### Computational Lesioning of Brain Nodes Predicts Callous-Unemotional Traits

Feature selection identified 23 brain nodes, modularity, and Tanner stage as features that improved prediction of CU traits (see Table 1 for coordinates of brain features and Figure 1C for all features). The elastic net model performed better than a dummy model and we found no evidence of overfitting (Table 2). As shown in Figure 1A, the association between predicted score and observed score for CU traits was significant (R^2^ = 0.311, p_perm_<0.001). The observed R^2^ in the permutation distribution is plotted in Figure 1B. Figure 1C plots distribution of cross validation betas for each feature in the model. These results suggest that the elastic net model performed satisfactorily in predicting CU traits.

**Figure 1.**
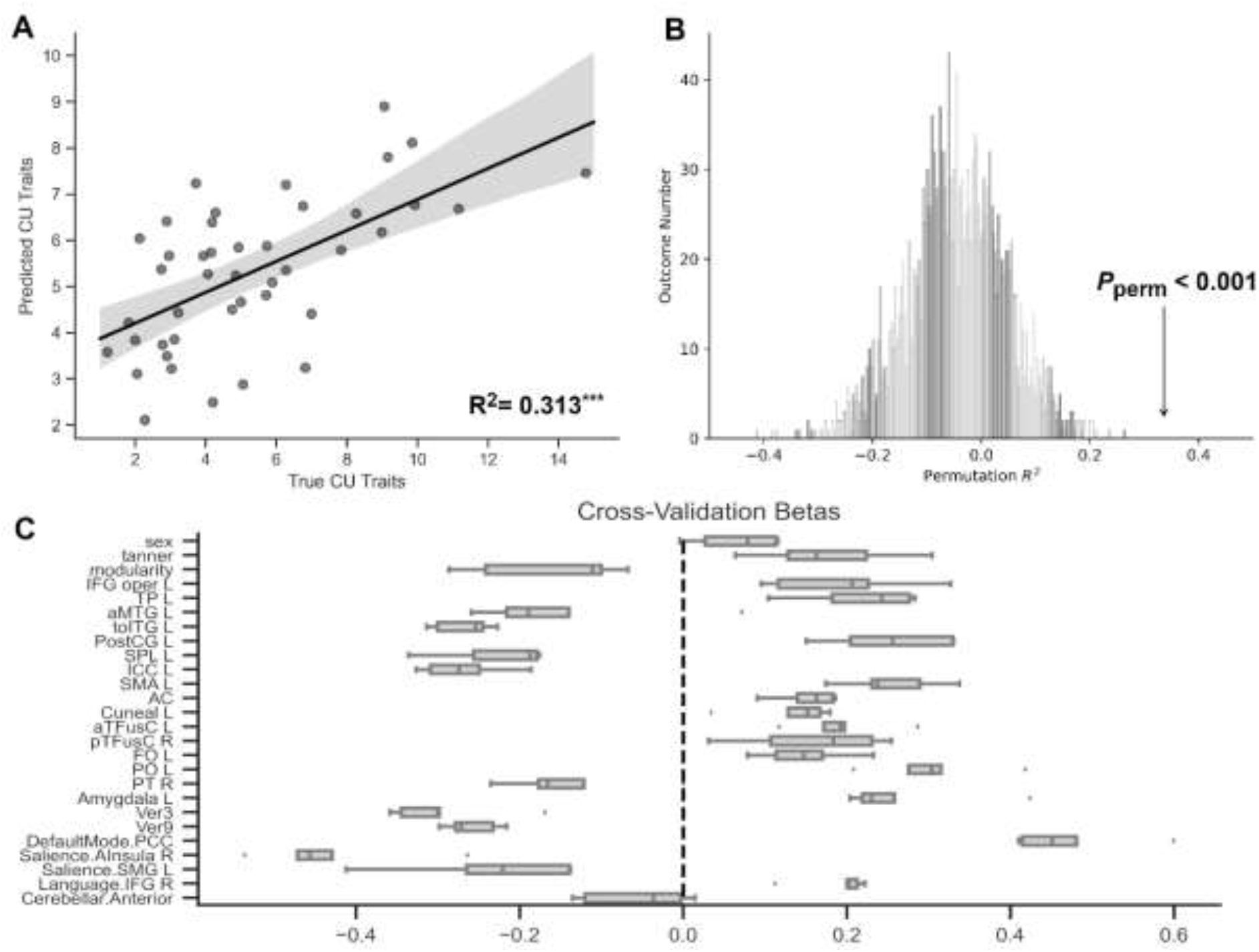
Predicting callous-unemotional traits with elastic net regression using selected features. A) Regression line and true R^2^ value. B) 2000 permutations testing of R^2^ values for the model and permuted p-value for true R^2^ value. C) 5-fold cross validation betas for each feature in the model.

**Table 2.** Elastic Net Cross Validation Fit Statistics

| Comparison to a Dummy Model |  |  |  |
| --- | --- | --- | --- |
| Cross Validation | Mean Squared Error |  | Best Fit |
|  | Elastic Net | Dummy |  |
| 1 | 7.522 | 10.265 | Elastic Net |
| 2 | 4.168 | 5.946 | Elastic Net |
| 3 | 9.042 | 21.315 | Elastic Net |
| 4 | 5.875 | 7.941 | Elastic Net |
| 5 | 7.651 | 9.588 | Elastic Net |
| Elastic Net Model Fit |  |  |  |
|  | MSE | R<sup>2</sup> | MAE |
| Training model | 4.244±0.268 | 0.601±0.075 | 1.569±0.060 |
| Test model | 6.784±1.613 | 0.310±0.123 | 2.00±0.169 |
| Permutation R<sup>2</sup> |  |  |  |
|  | True R<sup>2</sup> | Distribution | P value |
| Permutation R<sup>2</sup> | 0.313 | 0.146±0.0922 | < 0.001 |

Ten features negatively associated with CU traits: modularity, aMTG L, toITG L, SPL L, ICC L, PT R, Ver 3 and 9, anterior insula R, and SMG L (all abbreviations are in Table 1). Modularity indicates those with higher CU traits tended to have a less modular structure. The brain nodes indicate that a greater decrement in efficiency after computationally lesioning that node associated with higher CU traits. Fourteen features positively correlated with CU traits: Tanner stage, IFG oper L, TP L, PostCG L, SMA L, AC, Cuneal L, aTFusC L, pTFusC R, FO L, PO L, Amygdala L, PCC, IFG R (all abbreviations are in Table 1). Tanner stage indicates that those further along in pubertal development demonstrate higher CU traits. The remaining nodes indicate that less of a change in efficiency after lesioning each node associated with more CU traits. The two features that could not be distinguished from zero were sex and the Anterior Cerebellar region. The full model weights for the features in the elastic net model and their corresponding R^2^ and p-values derived from permutations are provided in Table 3 and the brain region beta weights are depicted in Figure 2.

**Table 3.**
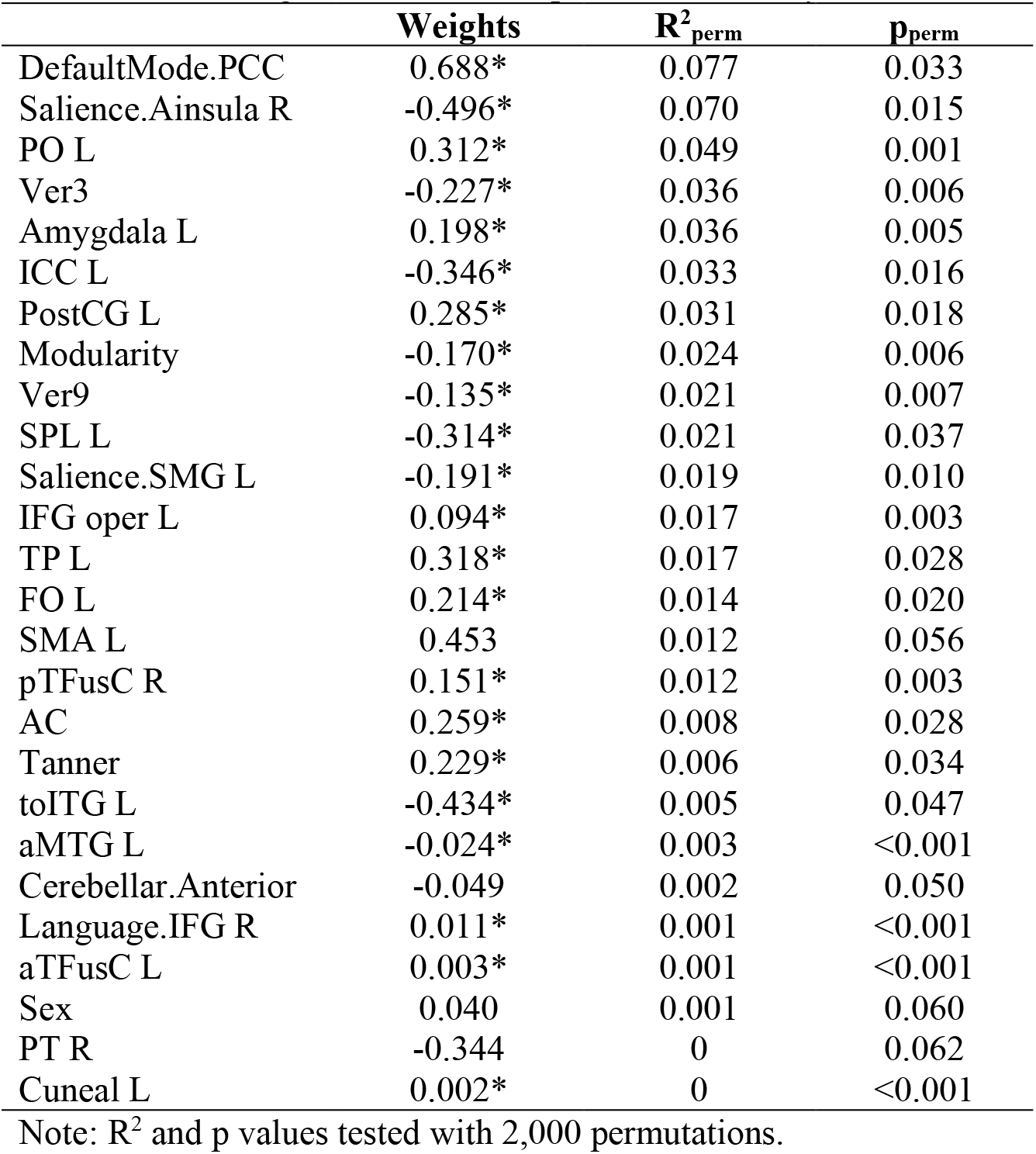
Beta Weights and Permuted p Values Sorted by Permuted R^2^

**Figure 2.**
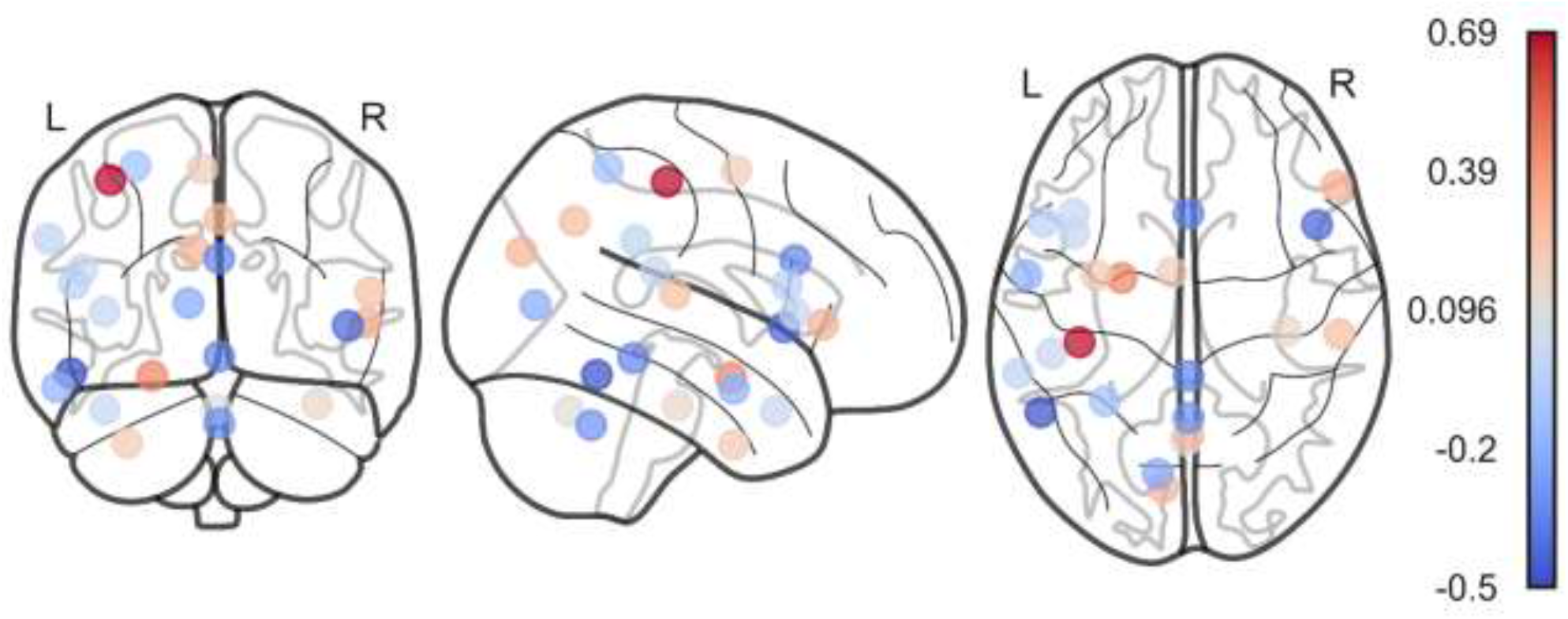
Brain region mask and corresponding beta values accounting for variance in callous-unemotional traits. Red and blue indicate positive and negative beta values, respectively. See Table 1 for coordinates of each region and Table 3 for specific beta values.

### Meta-Analytic Decoding of Region Mask Identified Emotional Terms

The mask of regions used in the elastic net model associated with several maps within Neurosynth covering emotions and affective information processing (Figure 3). The five terms with the highest associations involved the amygdala’s response to affective stimuli, mood, neutral (emotions), semantic control, and emotion regulation. When placed in the respective categories, emotional terms were of the most prominent category with 29 terms followed by 9 cognitive terms and 2 related to disorders (PTSD and anxiety disorder).

**Figure 3.**
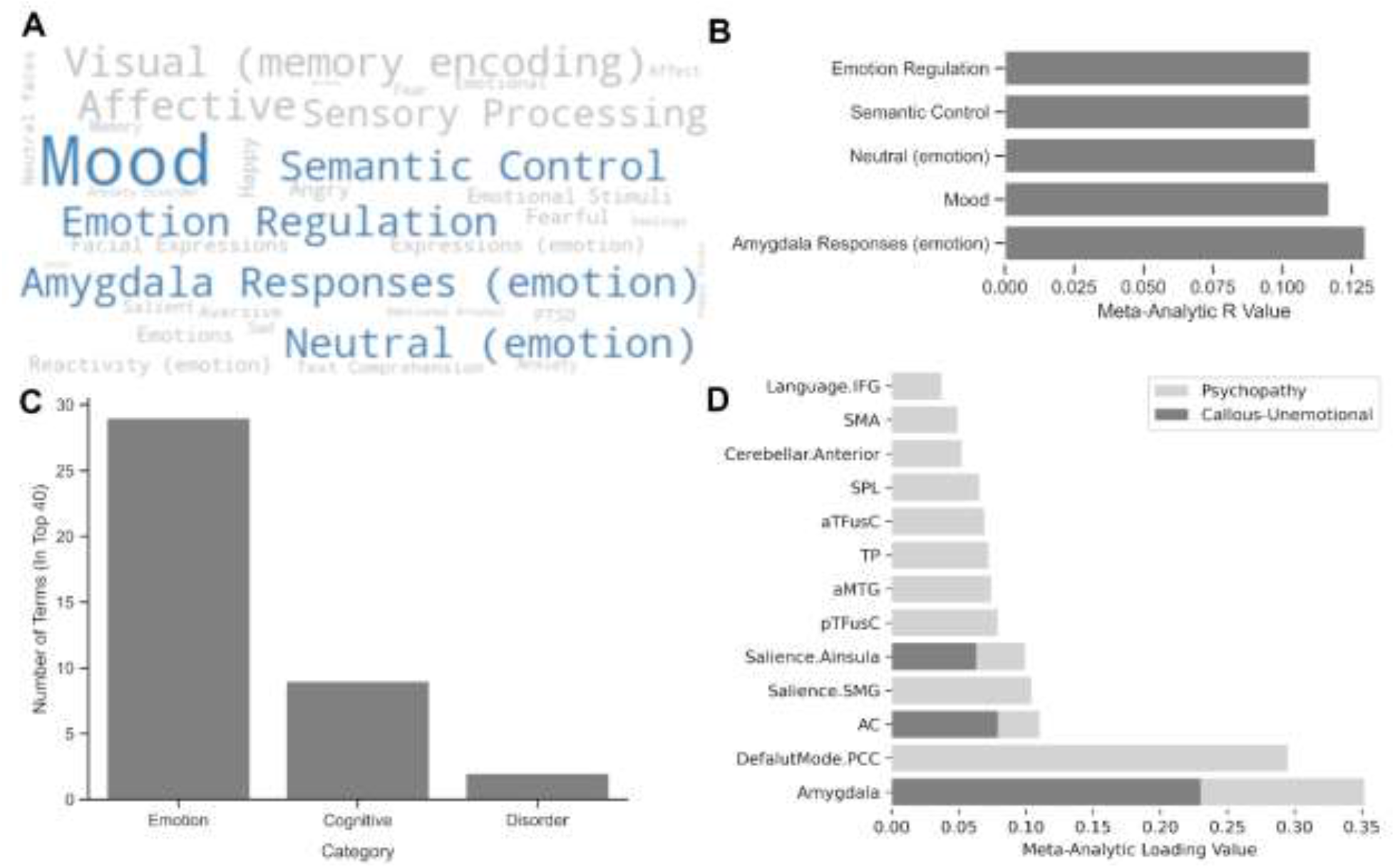
Meta-analytic decoding demonstrating that emotional terms associate with brain mask identified from simulated lesions and individual nodes loading on the callous-unemotional traits and psychopathy. A) Word cloud of top 40 terms where size indicates strength of correlation value and top five terms outlined in light blue; B) meta-analytic *r* values for the top five terms; C) categories of identified terms; D) loading of node terms on callous-unemotional traits or psychopathy.

### Meta-Analysis of Node Terms Loaded on Callous-Unemotional Traits and Psychopathy

A total of 13 nodes loaded on to CU traits and/or psychopathy, with the amygdala having the highest loading value. Three of the nodes loaded on to CU traits and all 13 loaded on to psychopathy. A total of 10 nodes associated with adult psychopathy but had not yet been identified in youth with CU traits among the studies within Neurosynth.

### Node Hub Differences at Higher Callous-Unemotional Traits

There were only a few nodes for which hubness associated with CU traits, but these hubs were notably less likely to be a hub (either global or local) over the entire sample. For global hubs, only the IFG oper L being a global hub associated with higher CU traits (Std β = 0.178, p= 0.006). Importantly, across the entire sample the probability of that region being considered a hub was only 4.7% (Table 4) and was not better than chance at being a global hub. For local hubs, the aMTG L (Std β = 0.304, p= 0.002), SPL L (Std β = 0.287, p < 0.001), ICC L (Std β = 0.054, p= 0.025), Cuneal L (Std β = 0.110, p= 0.022), and IFG R (Std β = 0.078, p= 0.038) being a local hub associated with higher CU traits. Again, the probability of these regions being a local hub across the sample was low (< 17%) and not better than chance (Table 5).

**Table 4.**
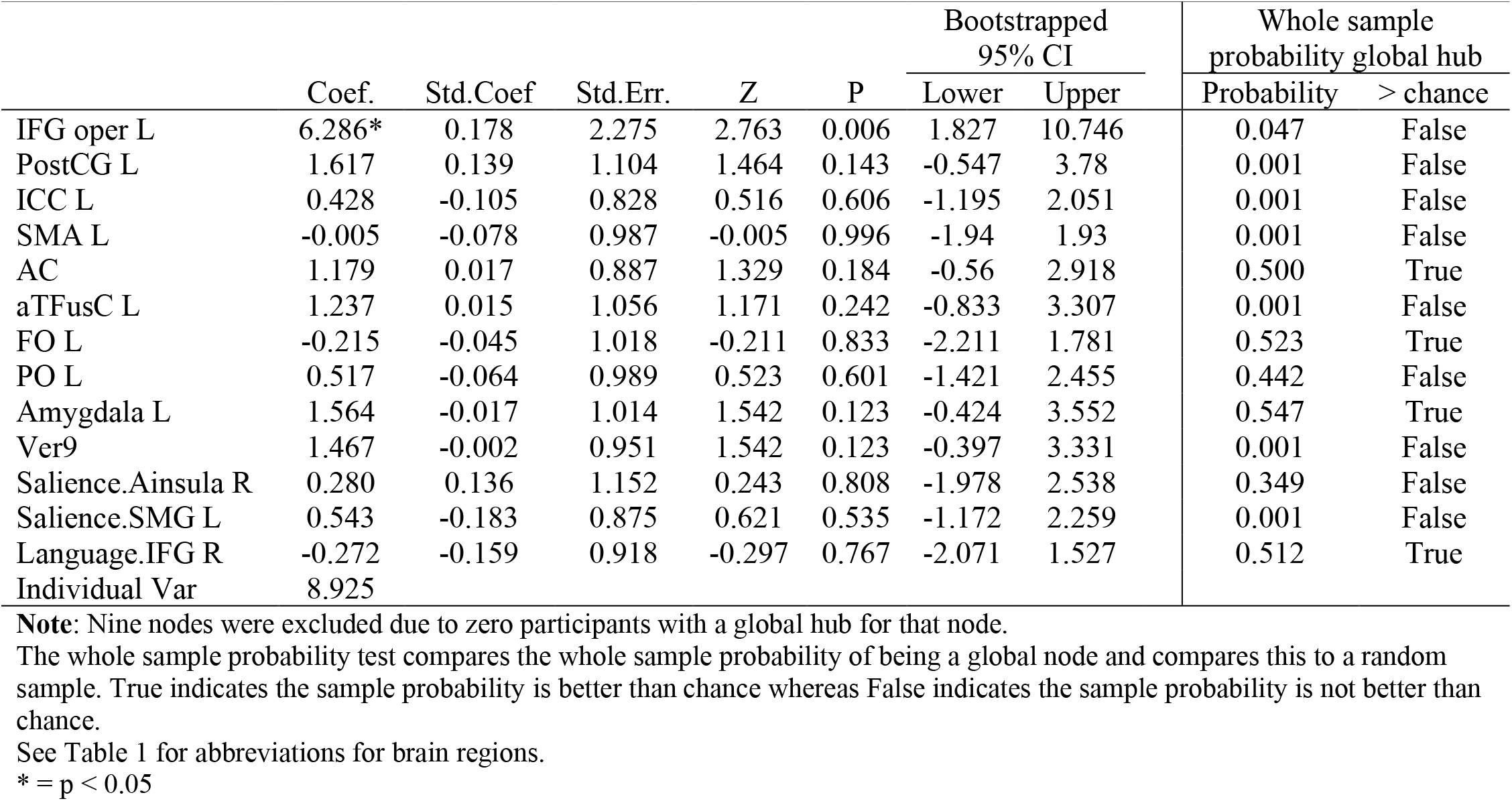
Results of Characterizing *Global* Nodes in Relation to Callous-Unemotional Traits

|  | Coef. | Std.Coef | Std.Err. | Z | P | Bootstrapped<br>95% CI |  | Whole sample<br>probability global hub |  |
| --- | --- | --- | --- | --- | --- | --- | --- | --- | --- |
|  |  |  |  |  |  | Lower | Upper | Probability | > chance |
| IFG oper L | 6.286* | 0.178 | 2.275 | 2.763 | 0.006 | 1.827 | 10.746 | 0.047 | False |
| PostCG L | 1.617 | 0.139 | 1.104 | 1.464 | 0.143 | -0.547 | 3.78 | 0.001 | False |
| ICC L | 0.428 | -0.105 | 0.828 | 0.516 | 0.606 | -1.195 | 2.051 | 0.001 | False |
| SMA L | -0.005 | -0.078 | 0.987 | -0.005 | 0.996 | -1.94 | 1.93 | 0.001 | False |
| AC | 1.179 | 0.017 | 0.887 | 1.329 | 0.184 | -0.56 | 2.918 | 0.500 | True |
| aTFusC L | 1.237 | 0.015 | 1.056 | 1.171 | 0.242 | -0.833 | 3.307 | 0.001 | False |
| FO L | -0.215 | -0.045 | 1.018 | -0.211 | 0.833 | -2.211 | 1.781 | 0.523 | True |
| PO L | 0.517 | -0.064 | 0.989 | 0.523 | 0.601 | -1.421 | 2.455 | 0.442 | False |
| Amygdala L | 1.564 | -0.017 | 1.014 | 1.542 | 0.123 | -0.424 | 3.552 | 0.547 | True |
| Ver9 | 1.467 | -0.002 | 0.951 | 1.542 | 0.123 | -0.397 | 3.331 | 0.001 | False |
| Salience.Ainsula R | 0.280 | 0.136 | 1.152 | 0.243 | 0.808 | -1.978 | 2.538 | 0.349 | False |
| Salience.SMG L | 0.543 | -0.183 | 0.875 | 0.621 | 0.535 | -1.172 | 2.259 | 0.001 | False |
| Language.IFG R | -0.272 | -0.159 | 0.918 | -0.297 | 0.767 | -2.071 | 1.527 | 0.512 | True |
| Individual Var | 8.925 |  |  |  |  |  |  |  |  |

**Note:** Nine nodes were excluded due to zero participants with a global hub for that node.
The whole sample probability test compares the whole sample probability of being a global node and compares this to a random sample. True indicates the sample probability is better than chance whereas False indicates the sample probability is not better than chance.
See Table 1 for abbreviations for brain regions.
\* = $p < 0.05$

**Table 5.**
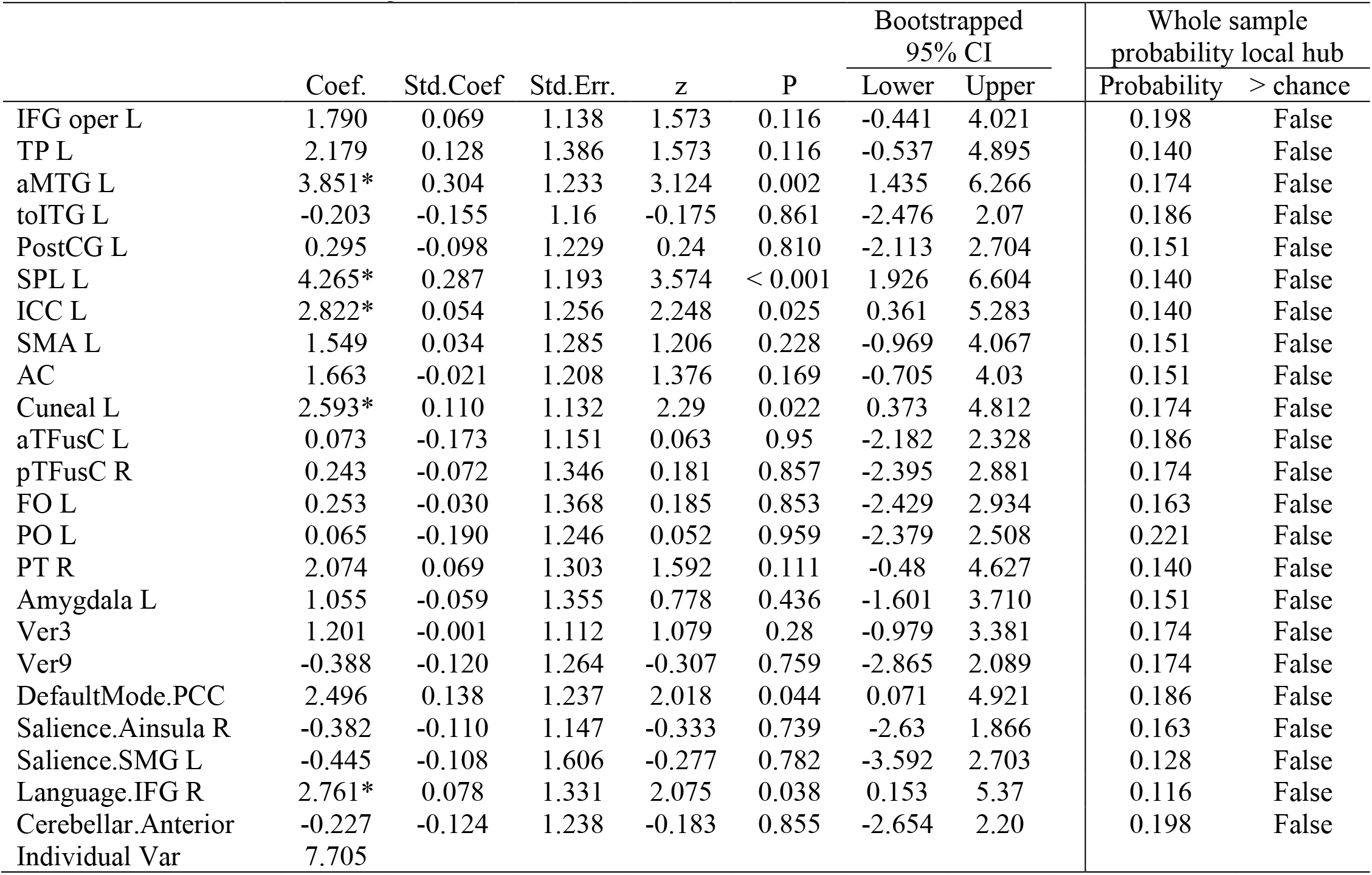
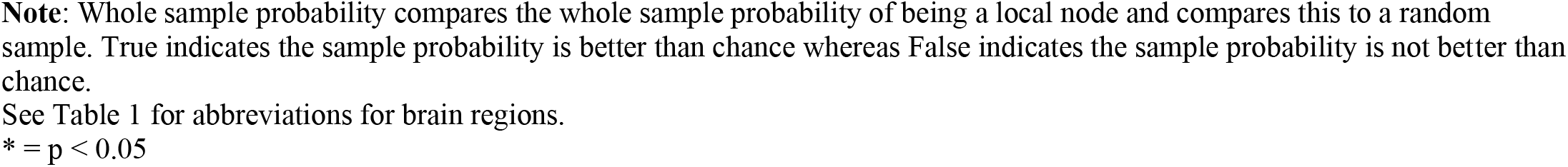
Results of Characterizing *Local* Nodes in Relation to Callous-Unemotional Traits

|  | Coef. | Std.Coef | Std.Err. | z | P | Bootstrapped<br>95% CI |  | Whole sample<br>probability local hub |  |
| --- | --- | --- | --- | --- | --- | --- | --- | --- | --- |
|  |  |  |  |  |  | Lower | Upper | Probability | > chance |
| IFG oper L | 1.790 | 0.069 | 1.138 | 1.573 | 0.116 | -0.441 | 4.021 | 0.198 | False |
| TP L | 2.179 | 0.128 | 1.386 | 1.573 | 0.116 | -0.537 | 4.895 | 0.140 | False |
| aMTG L | 3.851* | 0.304 | 1.233 | 3.124 | 0.002 | 1.435 | 6.266 | 0.174 | False |
| toITG L | -0.203 | -0.155 | 1.16 | -0.175 | 0.861 | -2.476 | 2.07 | 0.186 | False |
| PostCG L | 0.295 | -0.098 | 1.229 | 0.24 | 0.810 | -2.113 | 2.704 | 0.151 | False |
| SPL L | 4.265* | 0.287 | 1.193 | 3.574 | < 0.001 | 1.926 | 6.604 | 0.140 | False |
| ICC L | 2.822* | 0.054 | 1.256 | 2.248 | 0.025 | 0.361 | 5.283 | 0.140 | False |
| SMA L | 1.549 | 0.034 | 1.285 | 1.206 | 0.228 | -0.969 | 4.067 | 0.151 | False |
| AC | 1.663 | -0.021 | 1.208 | 1.376 | 0.169 | -0.705 | 4.03 | 0.151 | False |
| Cuneal L | 2.593* | 0.110 | 1.132 | 2.29 | 0.022 | 0.373 | 4.812 | 0.174 | False |
| aTFusC L | 0.073 | -0.173 | 1.151 | 0.063 | 0.95 | -2.182 | 2.328 | 0.186 | False |
| pTFusC R | 0.243 | -0.072 | 1.346 | 0.181 | 0.857 | -2.395 | 2.881 | 0.174 | False |
| FO L | 0.253 | -0.030 | 1.368 | 0.185 | 0.853 | -2.429 | 2.934 | 0.163 | False |
| PO L | 0.065 | -0.190 | 1.246 | 0.052 | 0.959 | -2.379 | 2.508 | 0.221 | False |
| PT R | 2.074 | 0.069 | 1.303 | 1.592 | 0.111 | -0.48 | 4.627 | 0.140 | False |
| Amygdala L | 1.055 | -0.059 | 1.355 | 0.778 | 0.436 | -1.601 | 3.710 | 0.151 | False |
| Ver3 | 1.201 | -0.001 | 1.112 | 1.079 | 0.28 | -0.979 | 3.381 | 0.174 | False |
| Ver9 | -0.388 | -0.120 | 1.264 | -0.307 | 0.759 | -2.865 | 2.089 | 0.174 | False |
| DefaultMode.PCC | 2.496 | 0.138 | 1.237 | 2.018 | 0.044 | 0.071 | 4.921 | 0.186 | False |
| Salience.Ainsula R | -0.382 | -0.110 | 1.147 | -0.333 | 0.739 | -2.63 | 1.866 | 0.163 | False |
| Salience.SMG L | -0.445 | -0.108 | 1.606 | -0.277 | 0.782 | -3.592 | 2.703 | 0.128 | False |
| Language.IFG R | 2.761* | 0.078 | 1.331 | 2.075 | 0.038 | 0.153 | 5.37 | 0.116 | False |
| Cerebellar.Anterior | -0.227 | -0.124 | 1.238 | -0.183 | 0.855 | -2.654 | 2.20 | 0.198 | False |
| Individual Var | 7.705 |  |  |  |  |  |  |  |  |

**Note:** Whole sample probability compares the whole sample probability of being a local node and compares this to a random sample. True indicates the sample probability is better than chance whereas False indicates the sample probability is not better than chance.
See Table 1 for abbreviations for brain regions.
\* = $p < 0.05$

### Targeting Global Hubs had Less Impact whereas Connector Non-Hubs Decreased Efficiency

Less change in efficiency after computationally lesioning global hubs positively associated with higher CU traits (Std β = 0.210, p= 0.006) but targeting local hubs did not produce significant effects (Table 6). A greater decrement in efficiency after lesioning connector non-hubs also associated with higher CU traits (Std β = –0.184, p= 0.024) but targeting peripheral nodes was not statistically significant (Table 7).

**Table 6.** Results of targeted lesions for global and local nodes predicting CU Traits

|  | Coef. | Std.Coef | Std.Err. | z | P | Bootstrapped<br>95% CI |  |
| --- | --- | --- | --- | --- | --- | --- | --- |
|  |  |  |  |  |  | Lower | Upper |
| Intercept | 6.468* | 0.000 | 1.173 | 5.514 | <0.001 | 1.678 | 11.771 |
| $\Delta$ Global Nodes | 20.641* | 0.210 | 7.567 | 2.728 | 0.006 | 1.714 | 42.429 |
| $\Delta$ Local Nodes | 30.113 | 0.140 | 17.183 | 1.753 | 0.08 | -12.041 | 71.427 |
| Modularity | -209.001* | -0.220 | 99.053 | -2.11 | 0.035 | -482.037 | 3.366 |
| Tanner | 0.812 | 0.237 | 0.355 | 2.285 | 0.022 | 0.062 | 1.509 |
| Sex | 0.078 | 0.012 | 0.715 | -0.109 | 0.913 | -1.752 | 1.560 |
| Individual Var | 4.997 |  |  |  |  |  |  |

Note: Global and local hubs were included in one analysis because of their statistical independence but not with their non-hub counterparts because of multicollinearity.
\* = $p < 0.05$

**Table 7.** Results of targeted lesions for connector and periphery nodes predicting CU Traits

|  | Coef. | Std.Coef | Std.Err. | z | P | Bootstrapped<br>95% CI |  |
| --- | --- | --- | --- | --- | --- | --- | --- |
|  |  |  |  |  |  | Lower | Upper |
| Intercept | -4.424 | -0.001 | 4.333 | -1.021 | 0.307 | -13.574 | 1.1781 |
| $\Delta$ Connector Nodes | -22.817* | -0.181 | 10.107 | -2.258 | 0.024 | -42.462 | -6.223 |
| $\Delta$ Periphery Nodes | -17.117 | -0.132 | 23.39 | -0.732 | 0.464 | -85.450 | 19.768 |
| Modularity | -204.058* | -0.215 | 99.714 | -2.046 | 0.041 | -490.497 | -18.660 |
| Tanner | 0.836* | 0.245 | 0.354 | 2.361 | 0.018 | 0.139 | 1.599 |
| Sex | 0.216 | -0.028 | 0.681 | 0.317 | 0.751 | -1.926 | 1.258 |
| Individual Var | 6.361 |  |  |  |  |  |  |

Note: Connector and periphery non-hubs were included in one analysis because of their statistical independence but not with their hub counterparts because of multicollinearity.
\* = $p < 0.05$

### No Evidence of Moderation

We found no statistically meaningful moderations for modularity or Tanner stage. Results of these analysis are placed in supplemental Tables 1 and 2.

### Heterogenous Functional Connectomes Evidence Weak Homogeneity

Figure 4 depicts the heterogeneity of individual brains even among those with higher or lower CU traits. Results were able to evidence some pattern level similarities at higher CU traits but there was considerable heterogeneity at the individual level.

**Figure 4.**
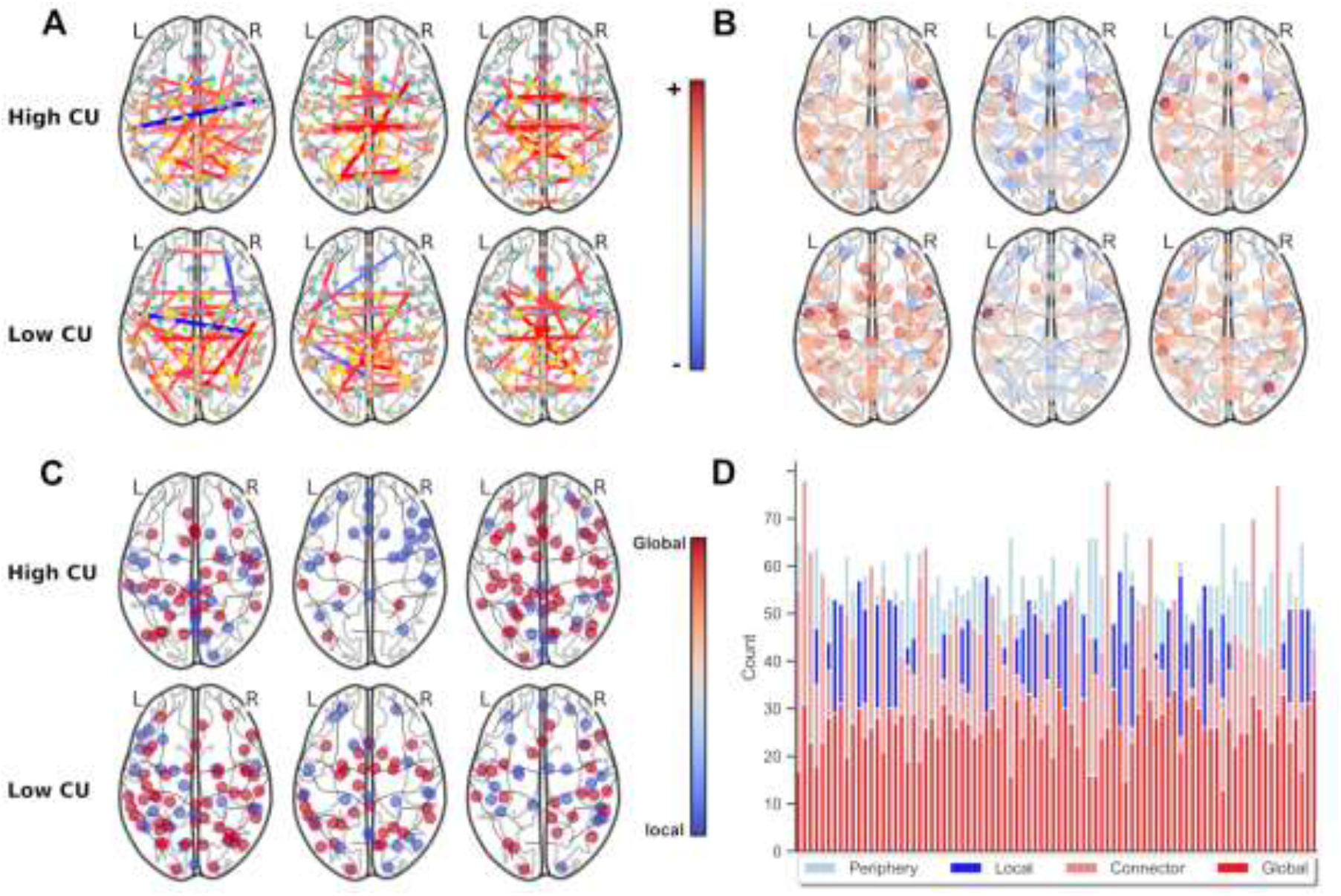
Adolescent brains are heterogenous even among those with similar scores at higher and lower CU traits. Three participants with high and low levels of CU traits were randomly selected to demonstrate this. A) individual-level functional connectomes; B) change in network efficiency after lesioning each node; C) location of global and local hubs; D) number of hubs and non-hubs for each individual participant, each column is a participant.

## Discussion

Overall results reveal that understanding efficiency changes after computational lesioning individual-level connectomes accounts for variation in CU traits in a community sample of adolescents. Moreover, the network properties of these nodes are important for distinguishing their impact on the network. The results of this study demonstrate the importance of topological features of the brain for gaining a mechanistic understanding of traits that may aid identification of individuals higher in CU traits.

### Lesioning Nodes Negatively Impacted Efficiency Across All Participants

As expected, the average response of the functional connectome for all selected nodes resulted in a reduction in efficiency. This reflects the importance of each individual node for the function of the entire brain, and removing any node has an impact on the brain’s efficiency. While this is a descriptive statistic, it is important for interpreting the results. Specifically, positive associations indicate resilience of the connectome because changes in efficiency are closer to zero whereas a negative association indicates a node where the functional connectome vulnerable.

### Computational Lesioning of Brain Nodes Predicts Callous-Unemotional Traits

As anticipated, feature selection identified cortical midline regions (e.g., anterior cingulate, posterior cingulate cortex) and regions associated with salience detection (right anterior insula, left supramarginal gyrus, and left anterior amygdala). What was less expected is that elastic net weights reveal important regions where the functional connectome is resilient where it was not expected to be. For example, the weight for the left amygdala was positive, suggesting that there was less of a change in efficiency after removing that node. The amygdala is a central hub for processing emotions and is heavily involved in social interactions (Bickart et al., 2014), thus we would expect greater decrements after its removal. However, the resilience of the connectome at higher CU traits after removing the amygdala suggests there is less integration with the rest of the connectome. Although many task-based studies found a lack of activation in the right amygdala at higher CU traits (Dotterer et al., 2017; Jones et al., 2009; Viding et al., 2012), our study, consistent with others (e.g., Marsh et al., 2008; Yang et al., 2009), reveals the left amygdala had less of an impact on the functional connectome’s efficiency. This discrepancy may be an artifact of stimuli presentation during a task-dependent state in prior studies (e.g., Funayama et al., 2001; Phelps et al., 2001). The present study did not impose a task-dependent state; thus, results plausibly reflect a more trait-like impairment involving less integration of the left amygdala associated with CU traits.

Additional positive associations were revealed in cortical midline structures including the posterior cingulate and anterior cingulate. The posterior cingulate is a central hub for information exchange (Leech et al., 2012) and is heavily involved in emotional arousal and attentional focus (Leech & Sharp, 2014) in healthy brains. The anterior cingulate is similarly involved in attention allocation, conflict monitoring, decision making, and emotion (Botvinick, 2007; Botvinick et al., 2004; Bush et al., 2000). As such, removal of these nodes was expected to impact the functional architecture of the brain in typically developing samples. However, less of a response of these nodes at higher CU traits suggests less integration with the rest of the connectome and less involvement in core processes for cognitive and emotional functioning.

However, lesioning other salience regions revealed a negative association of the right anterior insula and left supramarginal gyrus with CU traits. Where the right anterior insula is involved in attention (Eckert et al., 2009) involving interoceptive awareness (Craig, 2009, 2011) that aids both emotion and cognitive functioning (Touroutoglou et al., 2012), the left supramarginal gyrus is primarily involved in phonological processing (Celsis et al., 1999), though hypoactivity of this region has been associated with social cognition impairments in autism (Hadjikhani et al., 2006) and early psychosis (Park et al., 2021). The negative impact on the functional connectome after removing these nodes may indicate an overreliance on these regions for similar processes that would be distributed across the amygdala and other cortical midline structures, leading to decreased whole-brain global efficiency. The differences in the brains’ response to removal of these nodes characterize the impairments observed in CU traits as they appear to center around both cognitive and affective processing.

Modularity was another brain property accounting for variance in CU traits. Lower modularity, as revealed in prior work (Jiang et al., 2021), associates with higher CU traits. Beyond differences in specific connections, decreased modularity may better describe the apparent lack of convergence among functional connectivity studies of CU traits, all of which generally demonstrate less intra-network connectivity and abnormal inter-network connections (e.g., Cohn et al., 2015; Umbach & Tottenham, 2020; Winters, Sakai, et al., 2021; Yoder et al., 2016). This is an important property that may lead to a better understanding of multiple CU trait variants underlying individual differences (e.g., Fanti et al., 2013; Fanti et al., 2018; Hadjicharalambous & Fanti, 2018; Sebastian et al., 2012). Specifically, which networks and the extent to which they exhibit decreased modularity may underlie differences in CU trait profiles.

### Meta-Analytic Decoding of Mask Identified Emotional Terms

Meta-analytic decoding of the mask of regions identified in the present analysis primarily consisted of terms related to emotion and cognitive processes. Of the top 40 terms associated with the resulting mask, 29 were related to emotion processing (e.g., mood, amygdala response to emotion stimuli, and emotion regulation) and nine involved cognitive processes (e.g., semantic control, cognitive encoding, and memory). This constellation of associations is consistent with the broader literature and theoretical accounts that CU traits involve both cognitive and, more substantially, affective processing impairments.

### Meta-Analysis of Node Terms Loaded on Callous-Unemotional Traits and Psychopathy

Meta-analytic activations in the identified nodes revealed 13 individual nodes loading on either CU traits or psychopathy. Of these nodes, the five highest loading nodes were the amygdala, posterior cingulate, anterior cingulate, left supramarginal gyrus, and anterior insula. Importantly, ten of these nodes loaded only on psychopathy, which suggests our results revealed brain regions associated with a related adult phenotype that was not previously found in CU traits. It is therefore plausible that this mask may provide a comprehensive picture of neural underpinnings that could help predict CU traits.

### No Evidence of Moderation

There was no statistical evidence for modularity or Tanner stage moderating node efficiency changes related to CU traits. Moreover, the correlation between sex and CU traits did not meet study criteria to be further evaluated as a potential moderator. Overall, this suggests that the identified associations are direct and not affected by theoretically relevant, potential moderators.

### Node Hubness is Different at Higher Callous-Unemotional Traits

Both global and local hubs indicated at higher CU traits are not hubs in those that are lower in CU traits, suggesting a completely different topological structure at higher CU traits. Of the nodes surviving feature selection, a global hub in the IFG oper L and local hubs of the aMTG L, SPL L, ICC L, Cuneal L, and IFG R associated with higher CU traits, which was different from the rest of the sample. For example, the probability of the Inferior Frontal Gyrus Pars Opercularis L being a global hub across the entire sample was about 4.7% and not better than chance even though its “hubness” was associated with elevated levels of CU traits. Similar results were also found among the aforementioned local hubs. Conversely, the amygdala has been demonstrated to be an important hub in the brain (e.g., Bickart et al., 2014) and the full sample results support this finding (i.e., the left amygdala was identified as a global hub above chance). However, the left amygdala’s likelihood of being a hub was not associated with CU traits. This specificity of CU-related hubs further highlights the notion that certain patterns of variation in brain topology likely account for differences in the brains association with CU traits.

### Targeting Global Hubs Increased whereas Connector Non-Hubs Decreased Efficiency

Targeting hubs and non-hubs revealed some similarities but also important differences in functional architecture at higher CU traits. While targeting global hubs increased efficiency as expected, targeting local hubs had a positive trending yet non-significant effect despite predictions that doing so would decrease efficiency. Lesioning local hubs tends to remove shorter, more dense, and oftentimes more efficient connections, sparing longer and less efficient connections (Sporns et al., 2007; Tao & Rapp, 2021). The fact that functional connectivity patterns of those higher in CU traits correlate with less modularity – and are therefore less likely to have highly interconnected sets of local connections aiding efficiency – may explain why there was less of an impact on connectomes when attacking these nodes at higher CU traits.

Efficiency changes after computationally lesioning connector non-hubs associated negatively with CU traits. Because these nodes are more connected but do not possess the far reaching connections of global hubs (He et al., 2009; Tao & Rapp, 2021), these nodes are more likely to have shorter, more abundant connections that are critical for efficient functional connectivity. Thus, this finding is consistent expectations that lesioning connector non-hubs would logically negatively impact efficiency.

### Weak Homogeneity of Brains in Relation to Callous-Unemotional Traits

While pattern-level similarities were identified in relation to CU traits, substantial heterogeneity exists between individuals’ brains. This substantiates the weak homogeneity assumption (Gates, 2022) and, consistent with prior work (e.g., Dotterer et al., 2020; Winters, Sakai, et al., 2021), stresses the need to account for individual heterogeneity of the brains functional architecture in relation to CU traits. As such, we believe that our decision to not impose unrealistic homogeneity assumptions on adolescent brains lends itself to greater confidence in this study’s results.

### Limitations

The current study’s results should be interpreted with the following limitations in mind. First, the present study had a modest sample size that may have missed some important effects. For example, sex effects were trending as expected, but we may not have had sufficient power to detect these effects. Knowing whether sex moderates the relationships observed is worthy of future study, and the identified brain mask should be applied to larger samples with adequate distributions of males and females. Second, we sampled a range of ages that span multiple adolescent developmental stages (i.e., early to mid-adolescents), and larger samples within or across age bands would permit testing for age-specific differences. Third, while identified effects explained variance in CU traits, directionality cannot be determined from this cross-sectional sample. Longitudinal studies are needed to better examine the possibility of causality. Finally, the sample analyzed was a community sample that, although exhibiting similar neurocognitive (Viding & McCrory, 2012) and neurobiological impairments (Seara-Cardoso et al., 2022) as forensic individuals, may not adequately capture the extremely high levels of CU traits seen among forensic and clinical samples. As such, sampling community along with clinical and forensic samples are needed for future investigations of this nature.

### Conclusions

The present analyses demonstrate that knowing information about how individual-level brain connectomes respond to computationally simulated lesioning can explain meaningful variance in CU traits. This work identifies which nodes of adolescents’ functional connectomes are more resilient or vulnerable to computational lesioning and how this pattern differs among individuals with high levels of CU traits. For example, the left amygdala is a central and highly connected node that aids the coordination of salience and emotion processing in typically developing samples; the lack of response observed in the current study, however, suggests that the brain’s topological structure differs substantially at higher levels of CU traits. Similar results implicate the anterior cingulate and posterior cingulate cortices, cortical midline structures that are pivotal for conflict monitoring, attention, and emotion processing. These differences plausibly account for behavior differences related to neurocognitive and emotional processes observed among youth with elevated CU traits. Further topological differences were observed when lesioning local hubs did not impact efficiency as expected. These are important features describing qualities of information processing streams in the brain of youth with CU traits that suggest functional differences beyond activation or connection strength. Together, these results indicate a pattern of resilience and vulnerability, as well as underlying topological structure, characterizing CU traits. Future studies can advance our understanding of CU traits by using these identified regions to further investigate functional brain properties underlying CU traits as well as replication of predicting severity of CU traits.

## Supplemental material

**Supplemental Table 1.** Moderation results for Tanner

|  | Coef. | Std.Err. | z | P | Bootstrapped 95% CI |  |
| --- | --- | --- | --- | --- | --- | --- |
|  |  |  |  |  | Lower | Lower |
| aMTG.L.tanner | 964.836 | 657.374 | 1.468 | 0.142 | -323.593 | 2253.264 |
| ICC.L.tanner | -50.279 | 1041.484 | -0.048 | 0.961 | -2091.550 | 1990.993 |
| SMA.L.tanner | 651.300 | 1072.032 | 0.608 | 0.543 | -1449.844 | 2752.443 |
| aTFusC.L.tanner | 374.678 | 956.900 | 0.392 | 0.695 | -1500.812 | 2250.167 |
| Ver9.tanner | 300.325 | 769.044 | 0.391 | 0.696 | -1206.975 | 1807.624 |
| Salience.SMG.L.tanner | -959.656 | 1158.861 | -0.828 | 0.408 | -3230.982 | 1311.670 |
| aMTG L | -567.398 | 531.768 | -1.067 | 0.286 | -1609.645 | 474.849 |
| ICC L | -1114.639 | 839.558 | -1.328 | 0.184 | -2760.142 | 530.864 |
| SMA L | 2361.486 | 778.232 | 3.034 | 0.002 | 836.179 | 3886.792 |
| Ver9 | -1619.448 | 630.107 | -2.570 | 0.010 | -2854.435 | -384.460 |
| aTFusC L | 2058.055 | 785.255 | 2.621 | 0.009 | 518.984 | 3597.125 |
| Salience.SMG L | -1588.449 | 698.980 | -2.273 | 0.023 | -2958.425 | -218.474 |
| tanner | 0.452 | 0.347 | 1.303 | 0.193 | -0.228 | 1.133 |
| sex | 0.117 | 0.737 | 0.158 | 0.874 | -1.328 | 1.561 |
| modularity | -171.765 | 103.982 | -1.652 | 0.099 | -375.565 | 32.036 |
| Individual Var | 4.247 |  |  |  |  |  |

**Supplemental Table 2.**
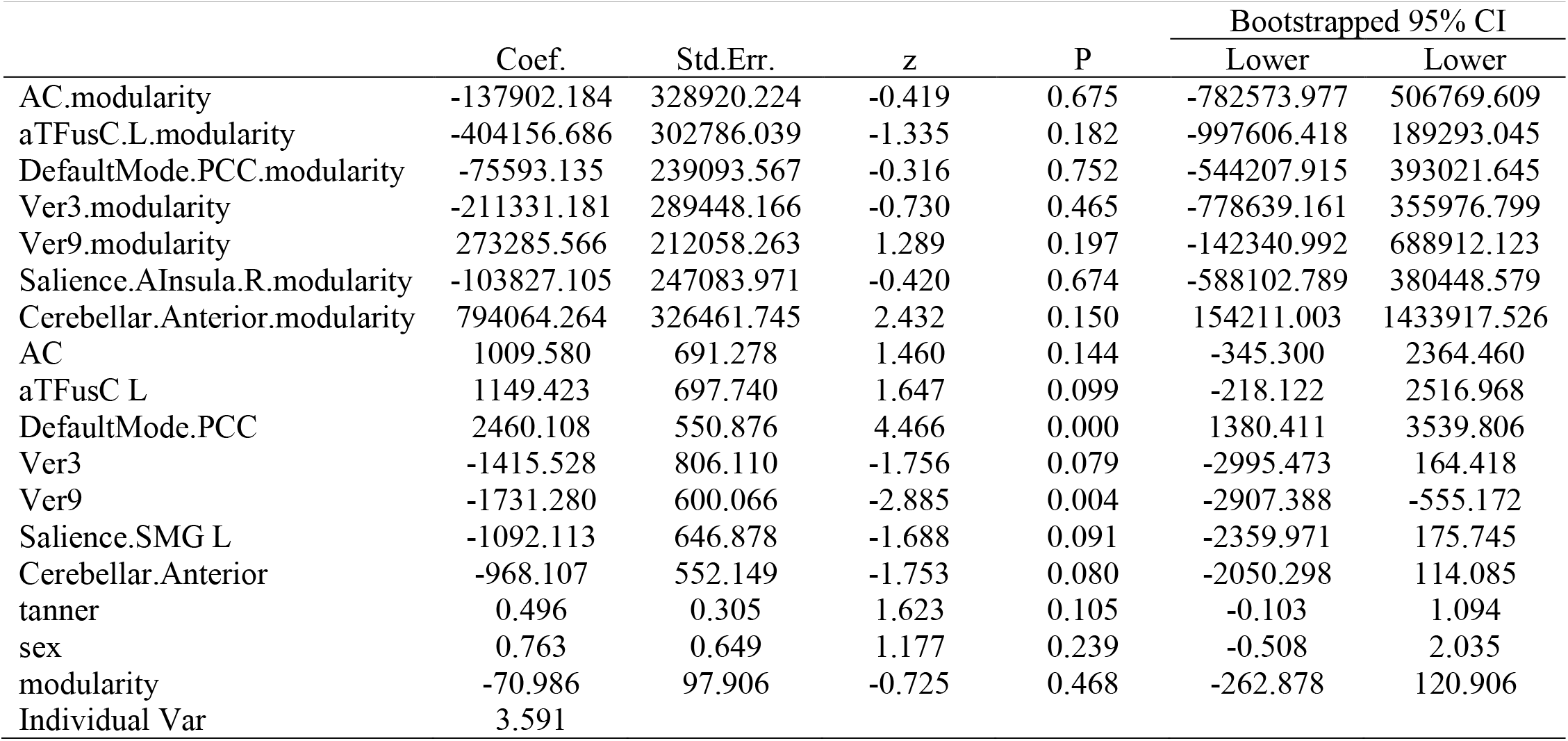
Moderation results for modularity

|  | Coef. | Std.Err. | z | P | Bootstrapped 95% CI |  |
| --- | --- | --- | --- | --- | --- | --- |
|  |  |  |  |  | Lower | Lower |
| AC.modularity | -137902.184 | 328920.224 | -0.419 | 0.675 | -782573.977 | 506769.609 |
| aTFusC.L.modularity | -404156.686 | 302786.039 | -1.335 | 0.182 | -997606.418 | 189293.045 |
| DefaultMode.PCC.modularity | -75593.135 | 239093.567 | -0.316 | 0.752 | -544207.915 | 393021.645 |
| Ver3.modularity | -211331.181 | 289448.166 | -0.730 | 0.465 | -778639.161 | 355976.799 |
| Ver9.modularity | 273285.566 | 212058.263 | 1.289 | 0.197 | -142340.992 | 688912.123 |
| Salience.AInsula.R.modularity | -103827.105 | 247083.971 | -0.420 | 0.674 | -588102.789 | 380448.579 |
| Cerebellar.Anterior.modularity | 794064.264 | 326461.745 | 2.432 | 0.150 | 154211.003 | 1433917.526 |
| AC | 1009.580 | 691.278 | 1.460 | 0.144 | -345.300 | 2364.460 |
| aTFusC L | 1149.423 | 697.740 | 1.647 | 0.099 | -218.122 | 2516.968 |
| DefaultMode.PCC | 2460.108 | 550.876 | 4.466 | 0.000 | 1380.411 | 3539.806 |
| Ver3 | -1415.528 | 806.110 | -1.756 | 0.079 | -2995.473 | 164.418 |
| Ver9 | -1731.280 | 600.066 | -2.885 | 0.004 | -2907.388 | -555.172 |
| Salience.SMG L | -1092.113 | 646.878 | -1.688 | 0.091 | -2359.971 | 175.745 |
| Cerebellar.Anterior | -968.107 | 552.149 | -1.753 | 0.080 | -2050.298 | 114.085 |
| tanner | 0.496 | 0.305 | 1.623 | 0.105 | -0.103 | 1.094 |
| sex | 0.763 | 0.649 | 1.177 | 0.239 | -0.508 | 2.035 |
| modularity | -70.986 | 97.906 | -0.725 | 0.468 | -262.878 | 120.906 |
| Individual Var | 3.591 |  |  |  |  |  |

